# Twenty-seven ZAD-ZNF genes of *Drosophila melanogaster* are orthologous to the embryo polarity determining mosquito gene *cucoid*

**DOI:** 10.1101/2022.09.04.506554

**Authors:** Muzi Li, Koray Kasan, Zinnia Saha, Yoseop Yoon, Urs Schmidt-Ott

## Abstract

The C2H2 zinc finger gene *cucoid* establishes anterior-posterior (AP) polarity in the early embryo of culicine mosquitoes. This gene is unrelated to genes that establish embryo polarity in other fly species (Diptera), such as the homeobox gene *bicoid*, which serves this function in the traditional model organism *Drosophila melanogaster*. The *cucoid* gene is a conserved single copy gene across lower dipterans but nothing is known about its function in other species, and its evolution in higher dipterans, including Drosophila, is unresolved. We found that *cucoid* is a member of the ZAD-containing C2H2 zinc finger (ZAD-ZNF) gene family and is orthologous to 27 of the 91 members of this family in *D. melanogaster*, including *M1BP, ranshi, ouib, nom, zaf1, odj, Nnk, trem, Zif*, and eighteen uncharacterized genes. Available knowledge of the functions of *cucoid* orthologs in *Drosophila melanogaster* suggest that the progenitor of this lineage specific expansion may played a role in regulating chromatin. We also describe many aspects of the gene duplication history of *cucoid* in the brachyceran lineage of *D. melanogaster*, thereby providing a framework for predicting potential redundancies among these genes in *D. melanogaster*.

## 1. Introduction

Dipteran insects (true flies) begin embryogenesis with 12 or 13 synchronous nuclear division cycles (1-4). During this syncytial phase of embryonic development, a uniform blastoderm forms in the cortical layer of the egg, activates the zygotic genome (5-7), and establishes axial polarity (8, 9). Anterior determinants (ADs) establish the embryo’s head-to-tail polarity via transcription factor gradients. In the fruit fly *Drosophila melanogaster* (*D. melanogaster*), the AD is encoded by the homeobox gene *bicoid* (10), which has been studied extensively (11-17). However, genes unrelated to *bicoid* are being used in species of different dipteran lineages for the same developmental task (18, 19). This evolutionary plasticity, along with the simple anatomy of early dipteran embryos and their amenability to experimental perturbation in non-traditional model organisms, set the stage for an attractive experimental system to study the molecular and evolutionary basis of transcriptional network stability and co-option of new central players in embryo development.

What are the molecular mechanisms that guide the co-option of new ADs? Bicoid has many target genes (15, 20-22), but it remains unclear how it adopted them. The *bicoid* gene evolved from a gene duplication of the *Hox3* ortholog of flies (also known as *zerknüllt* or *zen*), more than 145 million years ago (23-25). The diverged DNA-binding specificity of Bicoid, compared to its closest paralogs, prompted detailed studies on the evolution of its DNA-binding homeodomain using ancestral sequence reconstruction, quantitative *in vitro* DNA binding assays, and *in vivo* rescue experiments in Drosophila embryos (26-29). These studies emphasized the importance of mutations that altered DNA-binding specificity of the Bicoid protein. It was also shown that a feed-forward relay integrates certain regulatory activities of Bicoid and Orthodenticle via shared DNA binding sites (28). These homeodomain proteins have qualitatively similar DNA affinity and Orthodenticle has a conserved zygotic function in head development, which raised the question of whether Bicoid took over functions of Orthodenticle (29). However, comparative studies revealed ADs with distinct DNA binding domains and DNA affinities and suggest that the AD of the last common ancestor of dipterans was encoded by *pangolin* (*Tcf*) (18). Therefore, the co-option of new ADs in different fly lineages may not require shared target sites between the old and new ADs.

Why do specific genes adopt the AD function in addition to their other roles? The identification of AD gene orthologs in *Drosophila melanogaster* provides a useful starting point because many of its gene functions have been analyzed. For example, *odd-paired*, the only *zic* (*zinc finger of the cerebellum*) gene family member of flies (30), opens specific chromatin regions to advance the temporal progression of zygotic pattern formation in Drosophila embryos (31, 32). This function appears to be conserved in moth flies, where *odd-paired* additionally adopted the AD function by acquiring a maternal transcription variant (18). The ability of Odd-paired protein to drive the accessibility of specific chromatin regions, which is also a property of Bicoid (22, 33), could have facilitated their convergent co-option as ADs.

In culicine mosquitoes (e.g., Aedes and Culex), a previously uncharacterized C2H2 zinc finger gene, named *cucoid*, adopted the AD function. In these species, three *cucoid* transcript isoforms of with alternative 3’ ends have been identified in embryos. The shortest isoform is expressed maternally and is localized at the anterior pole of the egg. In culicine mosquitoes, knockdown of *cucoid* by RNAi results in ectopic expression of posterior genes at the anterior and the double abdomen phenotype (18). However, the function of *cucoid* orthologs in other species is unknown and obscured by a complex evolution of this gene in higher flies, including th *Drosophila melanogaster* (18). Here we show that *cucoid* is a member of the ZAD-ZNF gene family and is orthologous to at least 27 of *D. melanogaster’s* 91 ZAD-ZNF genes. ZAD-ZNF gene family members encode C2H2 zinc finger proteins with an N-terminal Zinc-finger-associated domain (ZAD) (34-39). Most *cucoid* orthologs of *D. melanogaster* have not yet been characterized but those that have been named and studied predominantly function in early development and oogenesis and may affect chromatin states.

## 2. Materials and methods

### 2.1 Identification of *cucoid* orthologs

Cucoid orthologs in the gall midge *Contarinia nasturtii* (*C. nasturtii*) and non-dipteran insects were identified by reciprocal protein BLAST (https://blast.ncbi.nlm.nih.gov/Blast.cgi) (40) using as queries the full-length Cucoid sequences of *Culex quinquefasciatus* (C. *quinquefasciatus*; QFQ59547.1) and *Aedes agypti* (*A. agypti*; XP_021704552.1) (18). Candidate orthologs were examined by conserved domain search against the Pfam database (41) (for presence of the ZAD) (https://www.ncbi.nlm.nih.gov/Structure/bwrpsb/bwrpsb.cgi) (42) using an E value cut-off of ≤ 0.01.

To identify Cucoid orthologs in *D. melanogaster*, we performed protein BLAST with the previously identified 91 *D. melanogaster* ZAD-ZNF proteins in *Aedes aegypti* and *Culex quinquefasciatus* with an E value cut-off of ≤ 0.05. Sequences that find full-length Cucoid isoforms as the best hits were considered to be candidate Cucoid orthologs in *D. melanogaster*.

Conservation of *Drosophila melanogaster* genes in the Cucoid clade in *Hermetia illucens, Bactrocera dorsalis, Lucilia cuprina*, and *Drosophila virilis* was assessed by reciprocal protein BLAST. Syntenies of Cucoid orthologs in different species were examined in GenBank and illustrated using the IBS server (43). Accession numbers are provided as supplementary material (**S1 Table**).

Orthology in the robber fly *Proctacanthus coquilletti* was established through tblastn (40) using *Hermetia illucens* and *D. melanogaster cucoid* orthologs as queries on the basis of query coverage and pairwise sequence identity. The identified orthologous exonic sequences were mapped to the robber fly genome (GCA_001932985.1), from which the full-length sequences were inferred **(S2 Table)**.

### 2.2 Phylogenetic analysis

The list of *D. melanogaster* ZAD-ZNF genes has been reported elsewhere (Kasinathan et al., 2020). The respective protein sequences were downloaded from GenBank. MAFFT alignments were generated by MAFFT v7.471with the L-INS-i strategy (https://mafft.cbrc.jp/alignment/software/) (44). For the protein tree with 91 ZAD-ZNF sequences, the raw alignments were trimmed using TrimAl v1.3 (http://phylemon2.bioinfo.cipf.es) (45) with a minimum percentage of positions to conserve of 20, a gap threshold of 0.8, and a similarity threshold of 0.05, to remove highly variable positions (columns). The trimmed alignments were further divided into two partitions corresponding to ZAD and ZNF regions. The best molecular substitution model for each partition was selected by partition merging strategy (MFP+MERGE) using ModelFinder (46) implemented in IQ-TREE v2.1.3 (47), based on Bayesian Information Criterion (BIC). Maximum likelihood trees were then built based on the selected substitution models, with branch support values generated by the implemented ultrafast bootstrap approximation (48), setting replicates to 3000. A majority rule consensus tree was generated form bootstrap trees and visualized by FigTree v1.4.4 (http://tree.bio.ed.ac.uk/software/figtree/). Trees are unrooted unless otherwise stated. Accession numbers of all sequences used in protein trees are provided as supplementary material **(S1 Table)**.

## 3. Results and Discussion

### 3.1 Cucoid is a ZAD-ZNF protein with 27 orthologs in *Drosophila melanogaster*

The genome of *Drosophila melanogaster* encodes around 300 C2H2 zinc finger proteins (36, 38), including multiple candidate orthologs of *cucoid*. To aid in the identification of *cucoid* orthologs, we searched for diagnostic domains and motifs of Cucoid using protein alignments and protein folding software (49). The alignment was constructed with previously reported single-copy Cucoid orthologs from the mosquitoes *Culex quinquefasciatus, Aedes aegypti*, and *Anopheles gambiae* (Culicidae), the harlequin fly *Chironomus riparius* (Chironomidae), the moth fly *Clogmia albipunctata* (Psychodidae), and the crane fly *Nephrotoma suturalis* (Tipulidae) (18), and with single-copy Cucoid orthologs from the gall midge *Contarinia nasturtii* (Cecidomyiidae), the cat flea *Ctenocephalides felis* (Siphonaptera), and in the silk moth *Bombyx mori* (Lepidoptera) that we retrieved from sequences deposited in GenBank (**S3 Figure**). No *cucoid* orthologs were found in other insect orders, suggesting that *cucoid* evolved during the radiation of holometabolous insects.

Cucoid proteins typically contain five C2H2 zinc finger domains. However, the Cucoid ortholog of Chironomus lacks zinc fingers 4 and 5, and culicine mosquitoes also express shorter isoforms without the 5^th^ (Culex) or 5^th^, 4^th^, and C-terminal half of the 3^rd^ zinc finger domains (Aedes). We found that all these Cucoid orthologs also contain a conserved N-terminal domain, known as Zinc-finger-associated domain (ZAD; **Fig 1, S3 Figure**) (34, 37). The ZAD is stabilized by zinc coordination via four invariant cysteine residues and can drive dimerization (36, 50) and nuclear localization (51).

**Fig 1.**
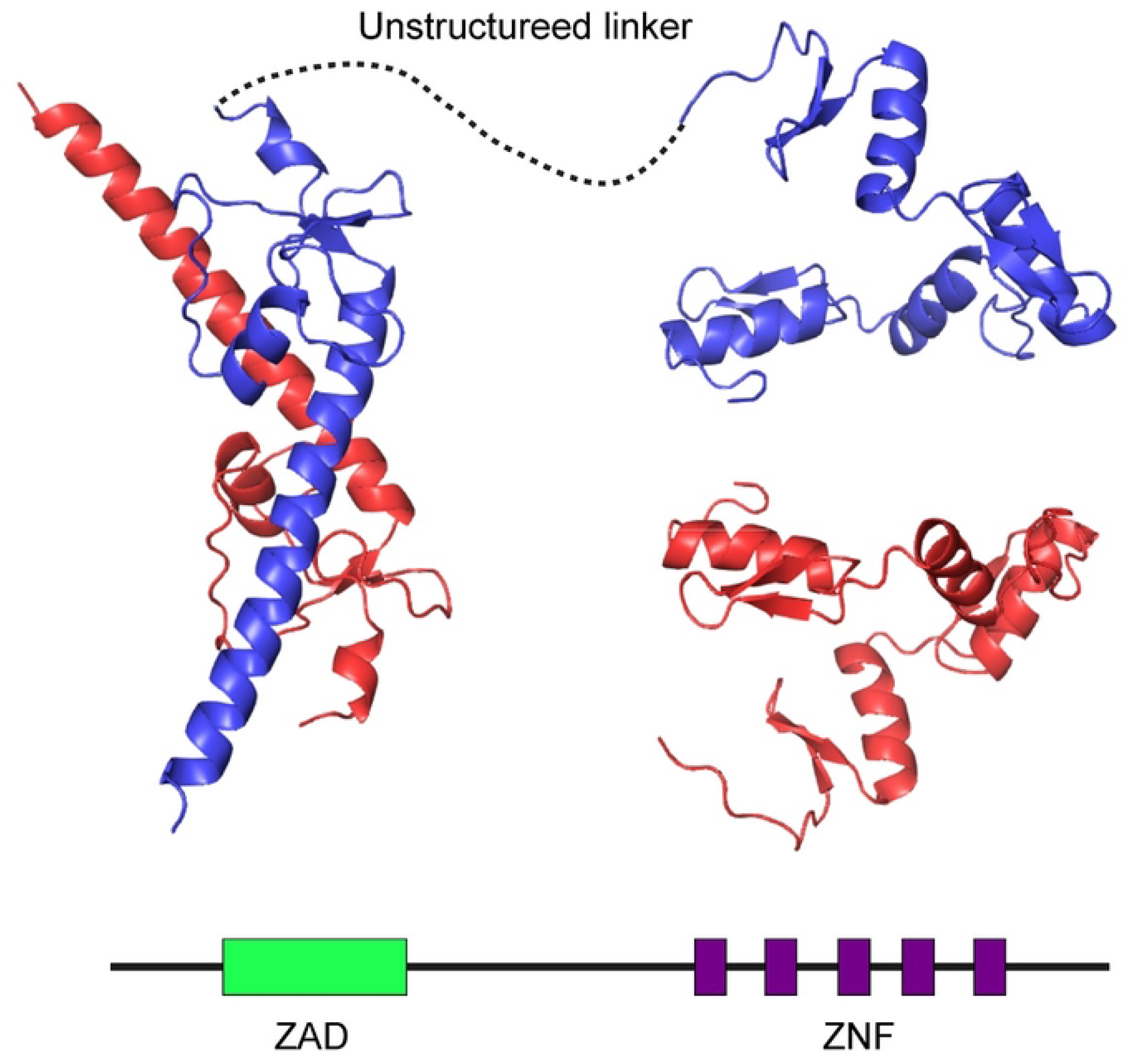
Predicted structure of Cucoid. Structure of a Cucoid homodimer (maternal isoform GenBank: QFQ59549.1) as predicted by Alphafold2 is shown with the two chains colored in red and blue above a simplified sketch of full-length Cucoid protein with the N-terminal end to the right and the C-terminal end to the left, and ZAD and ZNFs marked by colored rectangles.

Holometabolous insects evolved many ZAD-ZNF genes through lineage-specific gene duplications (34, 35, 38), especially in dipterans. For example, 147 ZAD-ZNF proteins have been found in *Anopheles gambiae* (35) and 91 in *Drosophila melanogaster* (39). To identify the ZAD-ZNF proteins in *D. melanogaster* most similar to Cucoid, we conducted protein BLAST with all 91 ZAD-ZNF proteins of *D. melanogaster* in *A. aegypti* and *C. quinquefasciatus*. The same seventeen Drosophila sequences retrieved *cucoid* in Aedes and Culex (54.1% sequence conservation). The corresponding genes were therefore considered candidate orthologs of *cucoid* (**Table 1**).

**Table 1.**
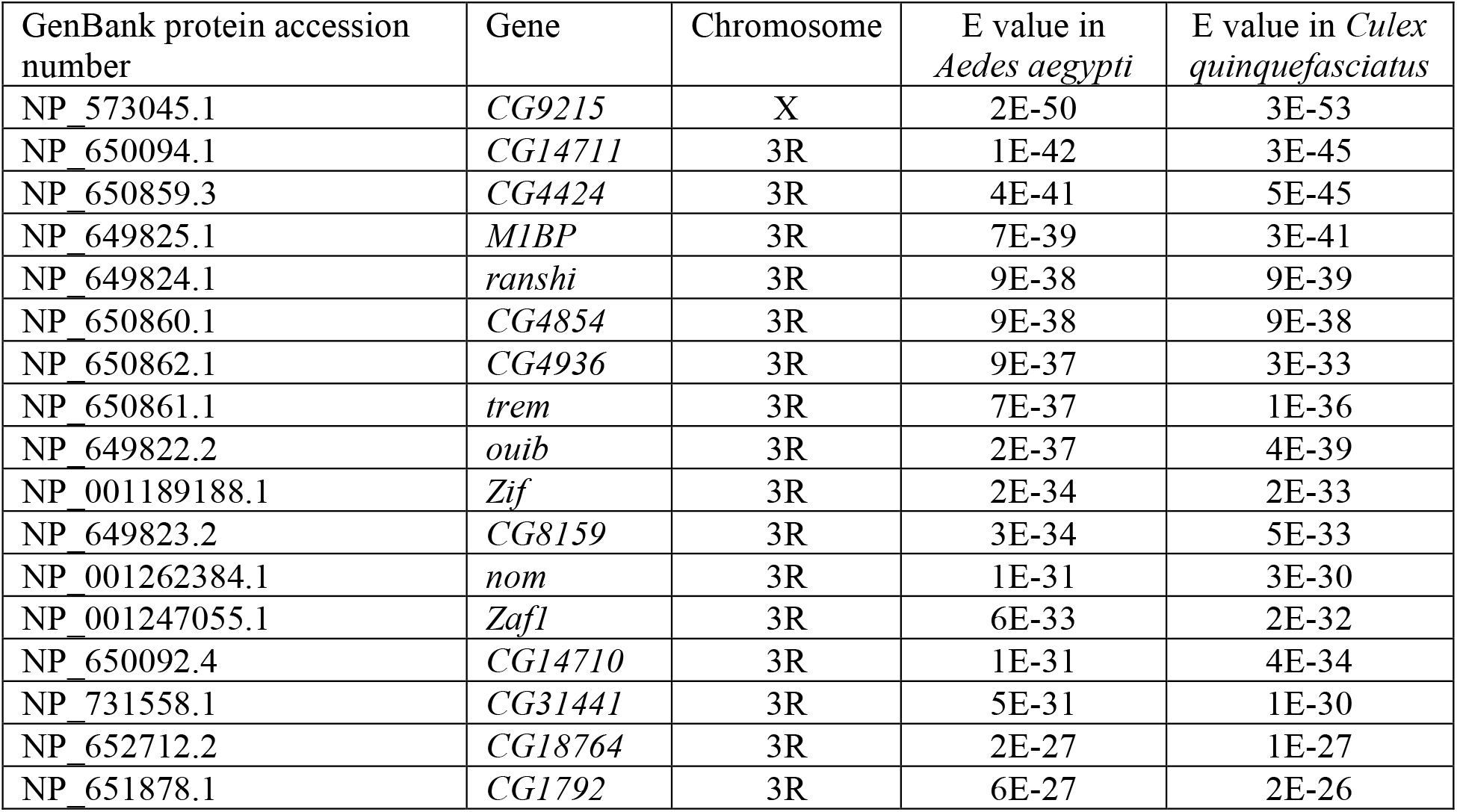
*D. melanogaster* ZAD-ZNF genes with *cucoid* as top BLAST hit in mosquitoes.

Next, we generated a Maximum Likelihood protein tree with all 91 ZAD-ZNF proteins of *D. melanogaster* and examined the distribution of the candidate Cucoid orthologs on this protein tree (**Fig 2**). All seventeen candidate orthologs (marked by red triangles in Fig. 2) mapped to a monophyletic clade of 27 ZAD-ZNF proteins (henceforth Cucoid clade). The Cucoid clade can be subdivided into two subclades with 9 members (subclade A) and 18 members (subclade B), respectively. All 9 members of subclade A were included in the original list of seventeen candidate Cucoid orthologs (**Table 1**). The top hit of that list, CG9215, also belongs to subclade A. The 18 members of subclade B experienced on average an elevated substitution rate. This subclade includes 10 genes that we did not recover using protein BLAST. These 10 genes do not form a monophyletic clade but are nested within the Cucoid clade and are therefore probably true Cucoid orthologs. We therefore conclude that at least 27 of *D. melanogaster*’s 91 ZAD-ZNF genes are orthologous to *cucoid*.

**Fig 2.**
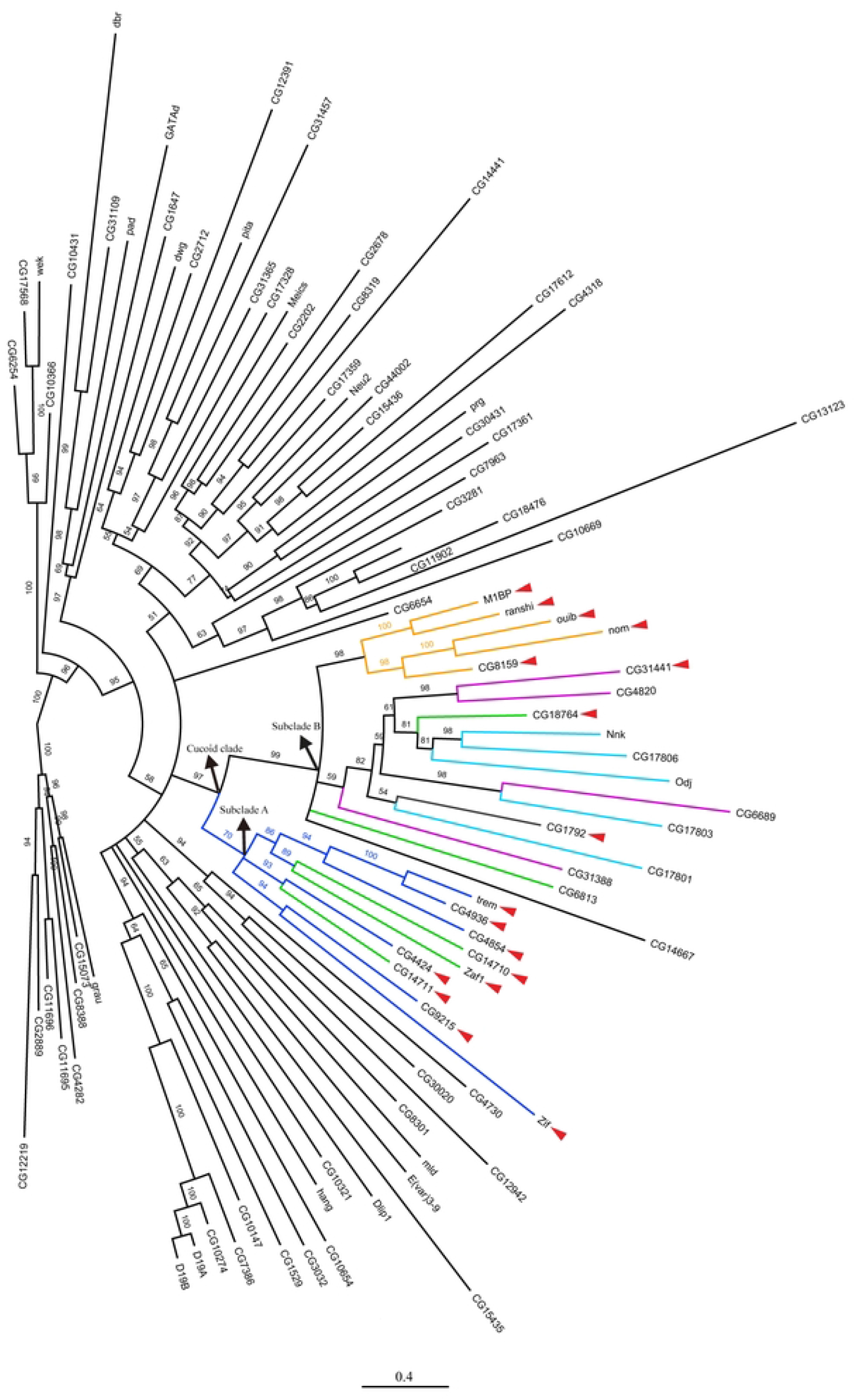
Manually rooted maximum likelihood protein tree of ZAD-ZNF family members in *D. melanogaster*. Orthologs that were identified based on reciprocal protein BLAST (see **Table 1**) are marked by red arrow heads. Colored lines correspond to chromosomal gene clusters (see **Fig 3**).

### 3.2 Genomic organization and relationship of genes of the Cucoid clade

All proteins of the Cucoid clade, except CG9215, are encoded by genes on the right arm of the 3^rd^ chromosome and are organized in five gene complexes of 4-5 genes and three isolated singletons (**Fig 3**). These 26 genes share intron positions among each other and with the *cucoid* orthologs from lower dipterans (**Fig 4**), consistent with an evolutionary origin by local gene duplications. *CG9215* is an intron-less gene on the X-chromosome that may have evolved by retro-transposition from the singleton *zif*, its putative sister gene (**Fig 2**). We denoted each gene cluster in *D. melanogaster* by the member that gave the smallest E value in reciprocal BLAST with Cucoid (**Table 1**), that is: M1BP cluster (orange), CG4424 cluster (blue), CG14711 cluster (green), and CG31441 cluster (purple). One cluster (light blue) did not include any of the genes that we identified by reciprocal BLAST and was named after a previously described gene, *oddjob* (*odj*) (52).

**Fig 3.**
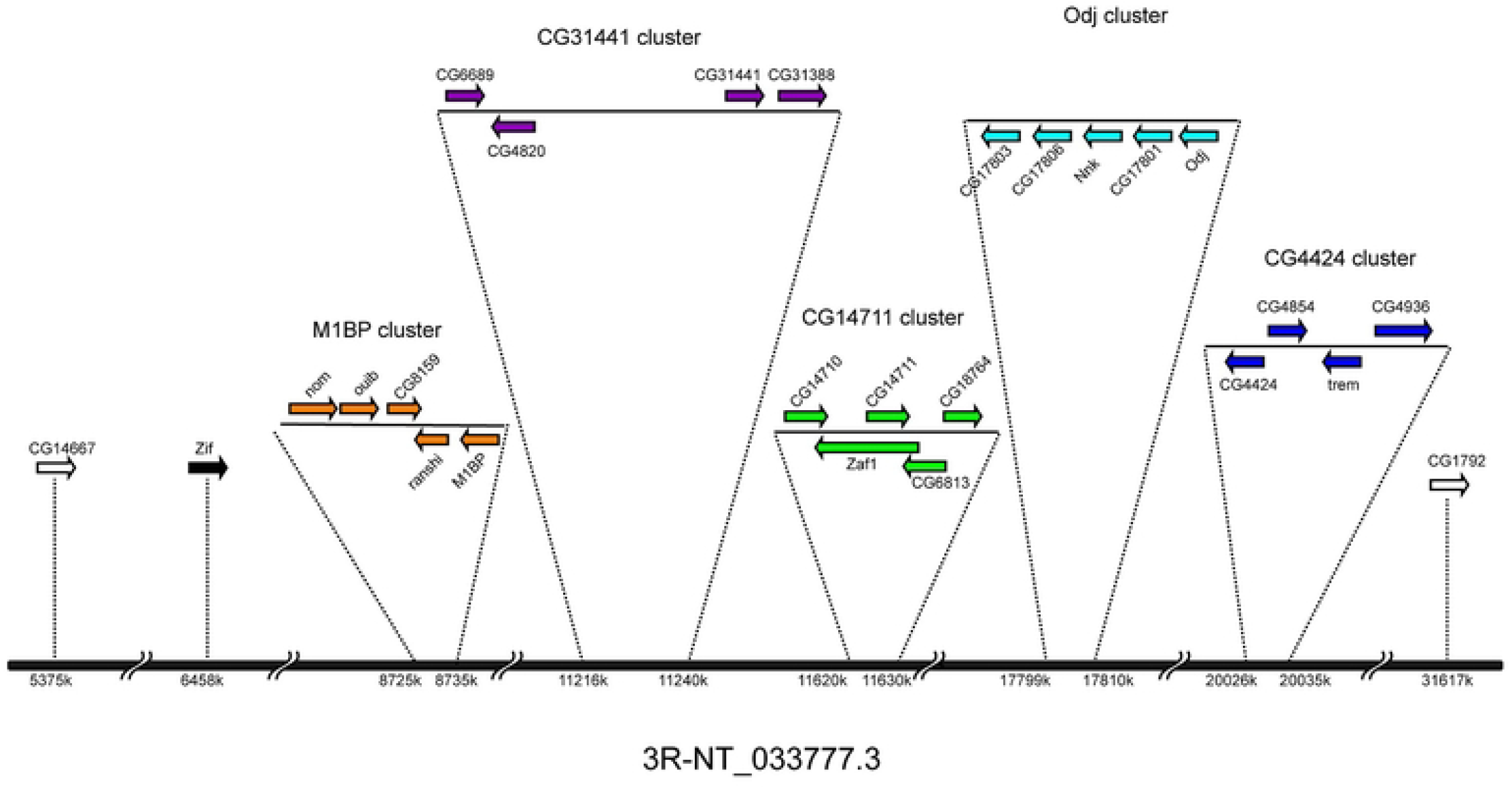
Synteny of genes in the Cucoid clade. Genes of the Cucoid clade are distributed in clusters on the right arm of the 3rd chromosome, except *CG9215* (not shown) which is located on the X chromosome. M1BP cluster (orange), CG14711 cluster (green), CG4424 cluster (blue), Odj cluster (light blue), CG31441 cluster (purple), *Zif* (black), other dispersed genes (outlined).

**Fig. 4.**
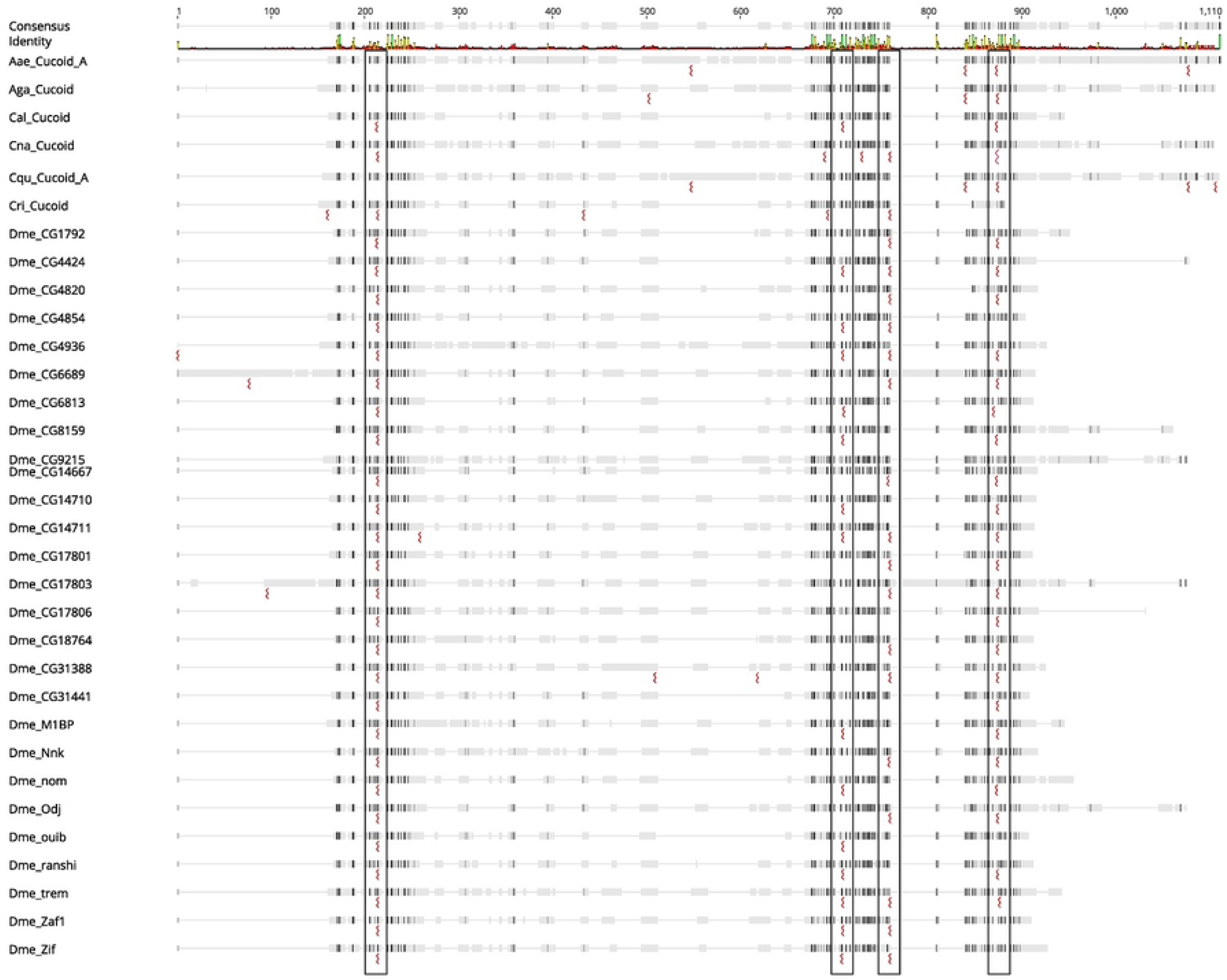
Conservation of Cucoid introns. Multiple sequence alignment of Cucoid orthologs from lower dipterans and Cucoid clade members of *D. melanogaster*. Conserved intron positions are boxed. Similarity of aligned amino acids is indicated by shading, with black representing high similarity and white representing no similarity. Sequence conservation and identity across all sequences are indicated in the top rows. Green, yellow, and red vertical lines indicate fully conserved, partially conserved, and poorly conserved amino acid residues, respectively.

The M1BP cluster genes form a monophyletic clade that can be traced to a single M1BP-like precursor gene, preserved in other schizophoran flies such as blow flies and tephritid fruit flies (see below). The first duplication of the *M1BP*-like precursor gave birth to *M1BP/ranshi* and *nom/ouib/CG8159*. The precursor of *nom/ouib/CG8159* duplicated twice, first generating *nom/ouib* and *CG8159* and then generating *nom* and *ouib* (**Fig 2**). These duplications occurred before the split of the *D. virilis* and *D. melanogaster* lineages (53). *M1PB/ranshi* duplicated after the split of *D. melanogaster* and *Drosophila ananassae* (53).

The other *cucoid*-related gene clusters of *D. melanogaster* do not form monophyletic clades. These incongruences between our protein tree and clustering in the *D. melanogaster* genome could have resulted from limitations of the phylogenetic inference methods that we used to build the protein tree, such as model choice and long-branch attraction (54), or non-allelic recombination within the ZAD-ZNF family (55, 56). However, in *D. virilis*, genes related to members of the CG14711 cluster (green), the CG31441 cluster (purple), and the Odj cluster (light blue) form a single gene complex with a different gene order, and this gene order is congruent with the inferred relationship of Cucoid clade genes (**Fig 5A**). Based on synteny in *D. virilis* and phylogenetic inference with *D. virilis* and *D. melanogaster* orthologs (**Fig 5B**), CG18764 of the CG14711 cluster (green) is sister to all genes of the Odj cluster (light blue) as well as CG6689 of the CG31441 cluster (purple). Additionally, we infer that CG6689 is sister to CG17803 of the Odj cluster, even though a gene-specific N-terminal THAP (Thanatos Associated Proteins) domain (57-59) that CG6689 inherited from its precursor, CG6689/CG17803, precursor is not preserved in CG17803 (**S4 Figure**). Finally, we infer that *D. virilis* lost the Nnk/CG17806 precursor because a CG17806-like precursor gene existed before the split of *D. melanogaster* and *D. virlis* but was not found in *D. virilis*. Duplication of the Nnk/CG17806 precursor occurred after the split of the *D. willistoni* lineage from *D. melanogaster* lineage (53).

**Fig 5.**
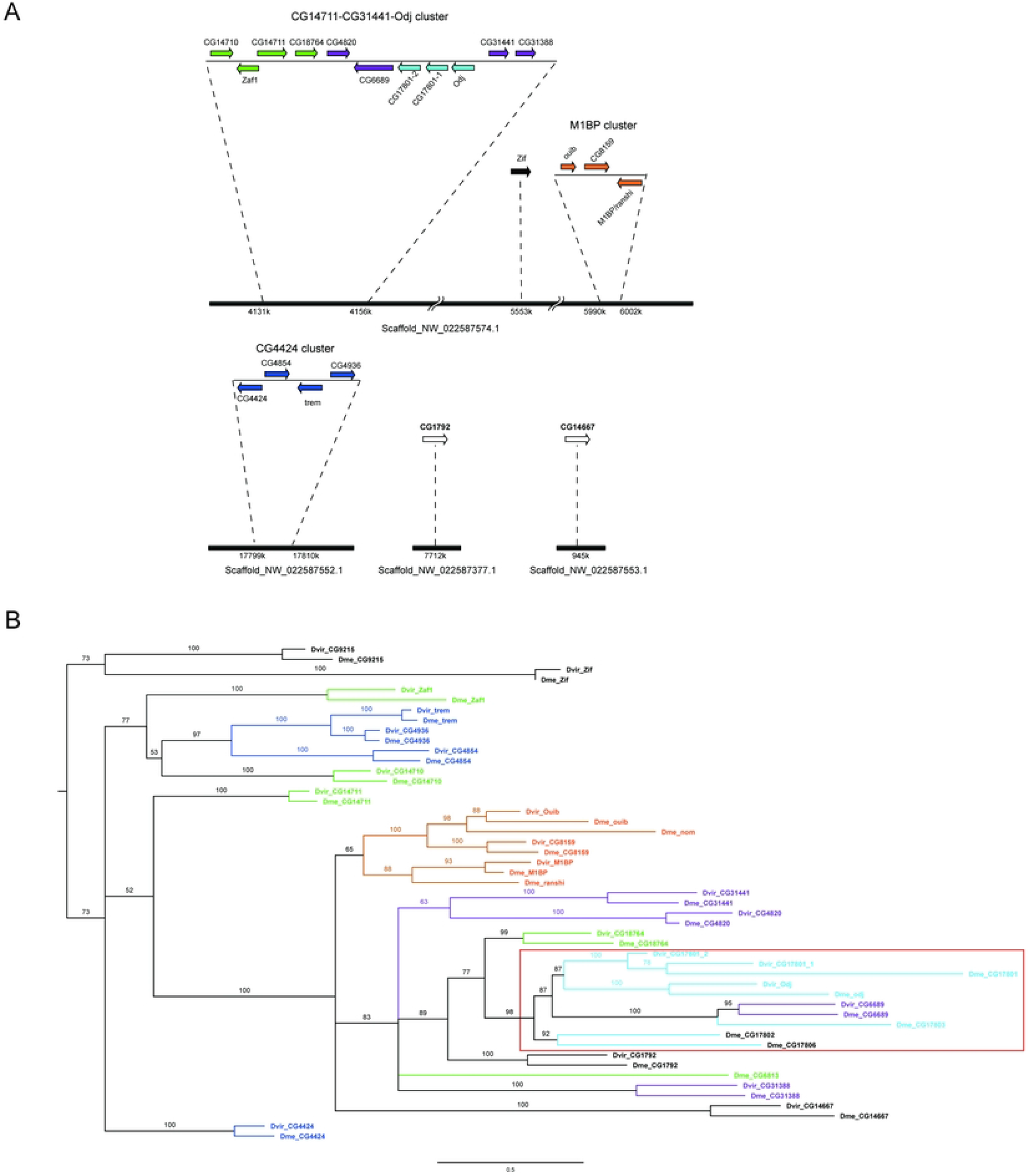
Cucoid clade protein tree is consistent with gene synteny in *D. virilis*. (A) Synteny of *cucoid* orthologs in *D. virilis*. Genes are color coded to indicate their relationship to gene clusters in *D. melanogaster* (see **Fig 3**). (B) Manually rooted maximum likelihood protein tree of Cucoid orthologs from *D. melanogaster* and *D. virilis*. Note that *D. virilis* has two CG17801 orthologs.

### 3.3 The Cucoid clade in Drosophila outgroups

Lineage-specific gene family expansions may reflect innovations or adaptations (37), but it is unknown why the number of ZAD-ZNF genes independently increased so much in multiple lineages of the Holometabola. To better understand when and how the Cucoid clade expanded, we searched for orthologs of the Cucoid clade members in representatives of other schizophoran fly species, including a blow fly (*Lucilia cuprina*) and a tephritid fruit fly (*Bactrocera dorsalis*), and in two representatives of the lower Brachycera, including a soldier fly (*Hermetia illucens*) and a robber fly (*Proctacanthus coquilleti*). While non-brachyceran dipterans have single *cucoid* orthologs (see section 3.1), we identified multiple *cucoid* orthologs in all the brachyceran species, albeit in lower numbers than in Drosophila.

In Lucilia and Bactrocera, we identified a single *M1BP*-like gene (orange), and several genes related to the CG4424 cluster (dark blue) and the CG14711 cluster (green), respectively, as well as putative orthologs of the sister genes *zif* and *CG9215* (**Fig 6A, B**). The presence of a *M1BP*-like gene in these unrelated species (they belong to paraphyletic lineages of the Schizophora (60)) suggests that the M1BP cluster expanded during, rather than before the radiation of the Schizophora in the Tertiary epoch (61). Whether the expansion of the M1BP cluster within the Schizophora resulted in subfunctionalization or the acquisition of new gene functions or a mix of both remains unknown, due to the lack of functional comparisons of the M1BP-like gene in lower Schizophora with their multiple orthologs in Drosophila.

**Fig 6.**
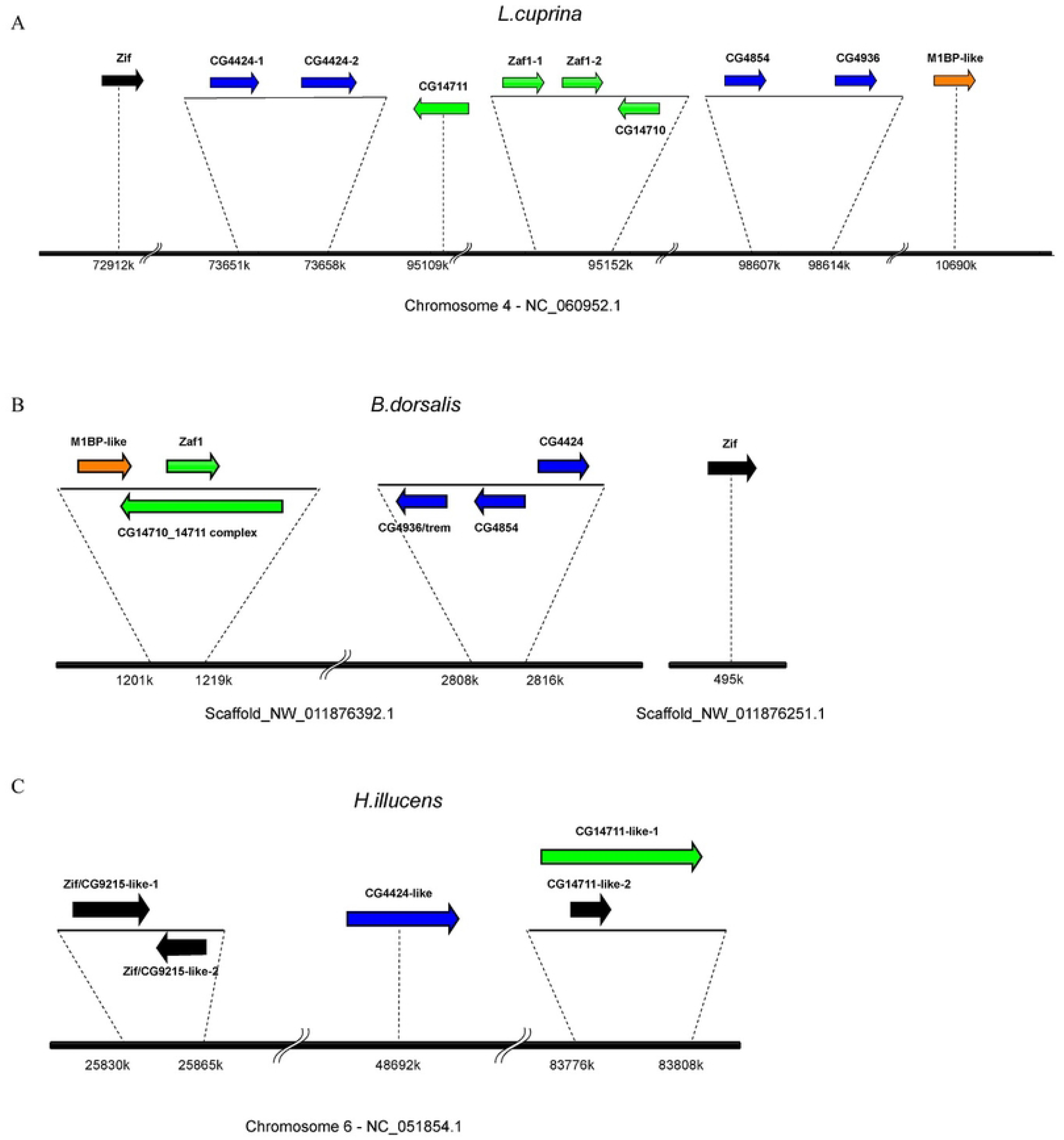
Syntenies of cucoid orthologs in Drosophila outgroups. Syntenies of cucoid orthologs in three brachyceran flies. Genes are color coded to indicate their relationship to gene clusters in *D. melanogaster* (A). Synteny of *cucoid* orthologs in *Lucilia cuprina*. A M1BP-like gene and three members in the CG14711 cluster (CG14711, CG14710, and Zaf1) are found, but other genes in *D. melanogaster*’s M1BP cluster were not present in this lineage. CG4424 cluster split into two loci, and two copies of CG4424 are in tandem. (B) Syteny of *D. melanogaster* cucoid orthologs conserved in *Bactrocera dorsalis*. M1BP locus is adjacent to CG14711 cluster, which may represent an ancestral state of these two clusters. (C) Genome locations of *D. melanogaster* cucoid orthologs in *Hermetia illucens. Hil*_1 and *Hil*_2 are paralogs born by tandem duplication and are proposed orthologous to CG9215/Zif. *Hil*_3 and *Hil*_4 are most closely related to CG4424 and CG14711 respectively by reciprocal BLAST. No *Hil_5* ortholog was found in *D. melanogaster* and the robber fly *Proctacanthus coquilletti*, suggesting it was born by lineage-specific duplication of its neighbor gene *Hil_4*. Thus, *Hil_5* was named by same orthologous relationship.

Two additional features of the genomic organization of Cucoid clade genes in *Bactrocera dorsalis* deserve attention. First, the *M1BP*-like gene of this species is in the immediate vicinity of the CG14711 cluster (green, **Fig 6A**). This finding suggests that the founder gene of the M1BP cluster originated as an offshoot of the CG14711 cluster, even though this is not apparent in the protein tree (**Fig 2**). Second, *CG14711* and *CG14710* of *B. dorsalis* have merged; the predicted protein has two linear ZAD-ZNFs structures that correspond to CG14710 and CG14711, respectively. Whether these genes are sister genes is unclear. Phylogenetic analysis suggests that the sister gene of *CG14711* is *CG4424* rather than *CG14710*. However, the predicted proteins of *CG14711* and *CG4424* are more Cucoid-like than the proteins of other members of the CG14711 cluster and the CG4424 cluster. Therefore, the inferred close relationship of CG14711 and CG4424 might reflect their less diverged status rather than their duplication history.

In the genomes of lower Brachycera, we identified five *cucoid* loci on chromosome 6 of the soldier fly *Hermetia illucens* (*Hil_cucoid_1-5*, GenBank accession numbers: XP_037925088.1, XP_037925165.1, XP_037924715.1, CAD7093451.1, XP_037922166.1) (62) (**Fig 6C**) and four *cucoid* loci in the robber fly *Proctacanthus coquilleti (Pco_cucoid_1-4)* (63) (**S2 Table**), which seem to be orthologous to Hermetia *cucoid* orthologs 1, 2, 3, and 4/5, respectively (**S5 Figure**). The lower brachyceran Cucoid proteins 1 and 2 are most similar to CG9215, but retain conserved introns and are therefore potentially orthologous to *CG9215/Zif*, whereas the lower brachyceran Cucoid orthologs 3 and 4 are most similar to CG4424 and CG14711, respectively. Therefore, the last common ancestor of soldier flies, robber flies, and Drosophila may have had at least three *cucoid* orthologs, including a *CG4424*-like member, *CG14711*-like member, and a CG9215-like ortholog of the Zif/CG9215 precursor.

## 4. Conclusions

*D. melanogaster* contains at least 27 *cucoid* orthologs, that is, almost one third of the 91 ZAD ZNF genes of this species. Reciprocal BLAST, phylogenetic inference, and genomic organization suggest that the Cucoid clade of *D. melanogaster* expanded gradually in the brachyceran lineage (**Fig 7**), while its founder gene was already present in the last common ancestor of butterflies, fleas, and flies. The last common ancestor of the brachyceran species that we analyzed may have had three *cucoid* orthologs that were similar to CG9215/zif, CG4424 and CG14711, respectively. The latter two may have been sister genes that generated all other genes of the Cucoid clade. The founder of the monophyletic M1BP-cluster originated before the radiation of the Schizophora. All other clusters of *D. melan*ogaster may not have monophyletic origins.

**Fig. 7.**
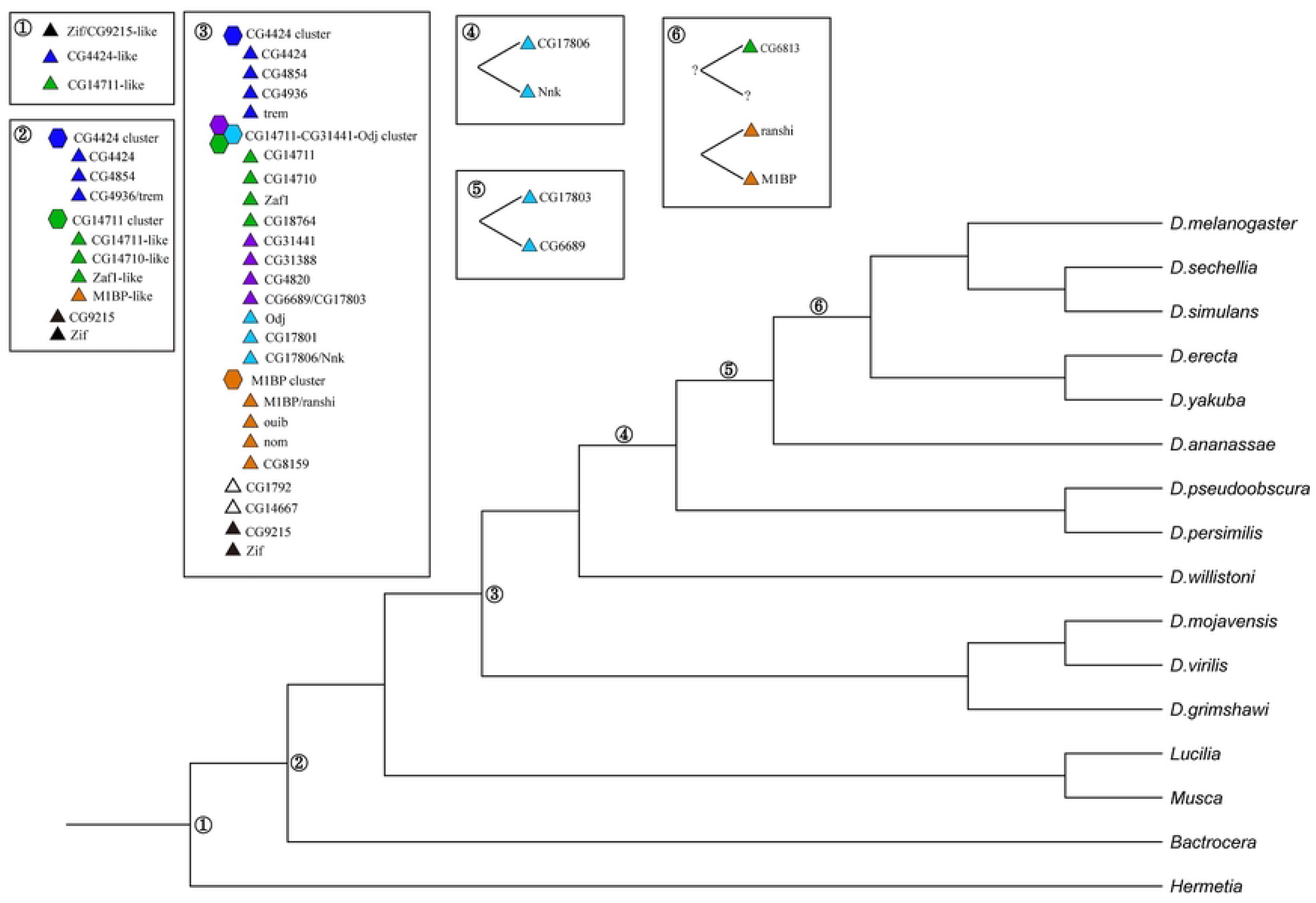
Evolution of the Cucoid clade family in higher flies. Inferred gene duplications in the Cucoid clade. For details and references see text. Gene cluster (hexagons) and gene loci (triangles) are indicated. For color code see Fig 2.

Our study was motivated by the question of what is known about *cucoid* orthologs in *Drosophila melanogaster*. Most of the 27 *cucoid* orthologs of *D. melanogaster* that we identified in this study did not affect viability when downregulated in previous large-scale screens (**Table 2**) (39, 64-66). However, several orthologs have been characterized in greater depth and display diverse, essential functions. For example, M1BP binds core promoters of thousands of genes and functions during transcription activation and polymerase II pausing while promoting chromatin accessibility surrounding the transcription start sites (67, 68). Other genes in the M1BP cluster show more specialized functions: *ranshi* regulates oocyte differentiation (69), *nom* functions in muscle development, and *ouib* is necessary during ecdysteroid synthesis by regulating *spookier* (70, 71). The closely related genes *odj* and *Nnk* have essential functions in heterochromatin regulation (39), *zaf1* is an architecture protein that serves as chromosome insulator in *Drosophila melanogaster* (72), *trem* is required for binding Mei-P22 on meiotic chromosomes to initiate double strand breaks for homologous recombination (73), and *Zif* is required for the expression and asymmetric localization of aPKC in neuroblast cells to regulate their polarity and self-renewal (74, 75). All other genes of the Cucoid clade remain uncharacterized. Taken together, our study suggests that many *cucoid* orthologs of *D. melanogaster* function in oogenesis and embryogenesis and several of them modify chromatin states. It will be interesting to find out whether the founder of the Cucoid clade also had a role in chromatin biology and whether Cucoid of culicine mosquitoes establishes embryo polarity epigenetically by regulating chromatin states of early zygotic segmentation genes in a concentration dependent manner.

**Table 2.** Functions of *D. melanogaster* cucoid orthologs.

| Gene | Functions | Loss-of-function | References |
| --- | --- | --- | --- |
| CG9215 | uncharacterized | viable | (64-66) |
| CG4424 | uncharacterized | viable | (64-66) |
| CG14711 | uncharacterized | unknown |  |
| Zaf1 | architectural protein,<br>chromosome insulator | viable | (64-66, 72, 76) |
| CG14710 | uncharacterized | viable | (39) |
| CG6813 | uncharacterized | viable | (39) |
| odj | heterochromatin binding | lethal | (39, 77) |
| ouib | regulates ecdysteroid synthesis;<br>stimulates transcription of<br>spookier | lethal | (70, 71) |
| M1BP | activates transcription;<br>orchestrates RNA polymerase II<br>pausing | viable | (67, 68) |
| Nnk | heterochromatin binding | lethal | (39, 64, 78) |
| ranshi | involves in oocyte<br>differentiation | sterile | (69) |
| CG17801 | uncharacterized | viable | (39, 66) |
| CG17806 | uncharacterized | viable | (39, 65) |
| CG17803 | uncharacterized | viable | (39, 64-66) |
| CG6689 | uncharacterized | lethal | (64) |
| CG31388 | uncharacterized | viable | (64, 65) |
| CG31441 | uncharacterized | lethal | (64) |
| CG4820 | uncharacterized | viable | (64-66) |
| CG18764 | uncharacterized | viable | (64-66) |
| CG1792 | uncharacterized | viable | (39, 65, 66) |
| Zif | regulates neuroblast<br>differentiation with *aPKC* | lethal | (64, 65, 74,<br>75) |
| CG14667 | uncharacterized | viable | (66) |
| CG8159 | uncharacterized | viable | (64) |
| nom | regulates embryonic muscle<br>morphogenesis | viable | (64-66, 79) |
| CG4936 | uncharacterized | lethal | (64, 65, 80) |
| trem | required for DSB formation in<br>meiosis | sterile | (73) |

## Acknowledgements

We thank our colleagues Dr. Phoebe Rice and Dr. Manyuan Long for helpful discussions, and Dr. Shengqian Xia and Dylan Sosa in the Long lab for technical assistance.

## Supporting Information

**S1 Table. Accession numbers**

**S2 Table. Locations of *cucoid* orthologs in *P. coquiletti* inferred from tblastn**

**S3 Figure. Cucoid protein alignment and prediction of Cucoid homodimer**

(A) Multiple sequences alignment of Cucoid orthologs from *Aedes aegypti* (Aae), *Anopheles gambiae* (Aga), *Bombyx mori* (Bmo), *Chironomus riparius* (Cri), *Clogmia albipunctata* (Cal), *Contarinia nasturtii* (Cna), *Ctenocephalides felis* (Cfe), *Culex quinquefasciatus* (Cqu), and Nsu (Nephrotoma suturalis). (B) A plot of the predicted alignment error of the best model acquired from Alphafold2 output which estimates the distance error for every pair of residues. Both axes represent the positions on the dimer of Cucoid maternal isoform (499 aa) from *C. quiquefasciatus*. The color key is measured in angstrom. Very low position errors are found for the overlapping of residues in the ZAD dimer as well as between zinc fingers on the same strand, indicating true packing of these domains. (C) The plot of predicted local distance difference test (pLDDT) per position gives a confidence level between 0-100 for each residue. All models predict ZAD and ZNF domain with very high confidence, whereas the highly variable linker regions get deficient support.

**S4 Figure. CG6689 acquired a DNA-binding THAP domain**

The THAP domains from 9 THAP-containing proteins in *D. melanogaster* and a THAP-like fragment from CG17803 are shown in alignment here. The color code for each column is based on similarity of aligned amino acids, with black representing high similarity and white representing no similarity. The THAP domain on the N-terminus of CG6689 is absent on other Cucoid orthologs including its sister gene CG17803 which has incomplete THAP features. THAP is a zinc-coordinating DNA binding domain with a conserved C2CH structure and shared features with the DNA binding domain of the P element transposase (57, 58). THAP-domain-containing proteins have been found in human, *D. melanogaster*, and *C. elegans* (57). In the 9 *D. melanogaster* proteins that have this domain, only CG6689 and CG10431 belong to the ZAD-ZNF family. CG10431 is only distantly related to CG6689 and located on a different chromosome (2L). The THAP domain on CG6689 is encoded by the first two exons of this gene. The intron mapping of *cucoid* orthologs **(Fig 4)** shows that the first exon of CG6689 is not present on other *cucoid* orthologs except for its sister gene CG17803 which have also lost the key residues defining the THAP domain. Homology search of its sequence by blastn only found hits within the Drosophila genus. Although it is unclear how THAP domain forms on unrelated proteins, the evidence here suggests a possibility of de novo formation.

**S5 Figure. Phylogeny of Cucoid orthologs in *H. illucnes* and *P. coquilletti***

A phylogenetic tree with Cucoid orthologs in *H. illucnes, P. coquilletti* and lower flies was constructed based on untrimmed alignment using 3 partitions inlucing ZAD, ZNF, and the left regions. Regions outside the conserved domains are preserved here to keep the diagnostic features useful for inferring orthlogy. From the branches with high support values, this tree suggest that *Hil_cucoid_1* to *Hil_cucoid_4* are orthologous to *Pco_cucoid_1* to *Pco_cucoid_4* respectively.

